# Microsecond simulations and CD spectroscopy reveals the intrinsically disordered nature of SARS-CoV-2 Spike-C-terminal cytoplasmic tail (residues 1242-1273) in isolation

**DOI:** 10.1101/2021.01.11.426227

**Authors:** Prateek Kumar, Taniya Bhardwaj, Neha Garg, Rajanish Giri

## Abstract

All available SARS-CoV-2 spike protein crystal and cryo-EM structures have shown missing electron densities for cytosolic C-terminal regions. Generally, the missing electron densities point towards the intrinsically disordered nature of the protein region. This curiosity has led us to investigate the C terminal cytosolic region of the spike glycoprotein of SARS-CoV-2 in isolation. The cytosolic regions is supposed to be from 1235-1273 residues or 1242-1273 residues depending on what prediction tool we use. Therefore, we have demonstrated the structural conformation of cytosolic region and its dynamics through computer simulations up to microsecond timescale using OPLS and CHARMM forcefields. The simulations have revealed the unstructured conformation of cytosolic region. Also, in temperature dependent replica-exchange molecular dynamics simulations it has shown its conformational dynamics. Further, we have validated our computational observations with circular dichroism (CD) spectroscopy-based experiments and found its signature spectra at 198 nm which is also adding the analysis as its intrinsically disordered nature. We believe that our findings will surely help us understand the structure-function relationship of the spike protein’s cytosolic region.

**Graphical Abstract:** 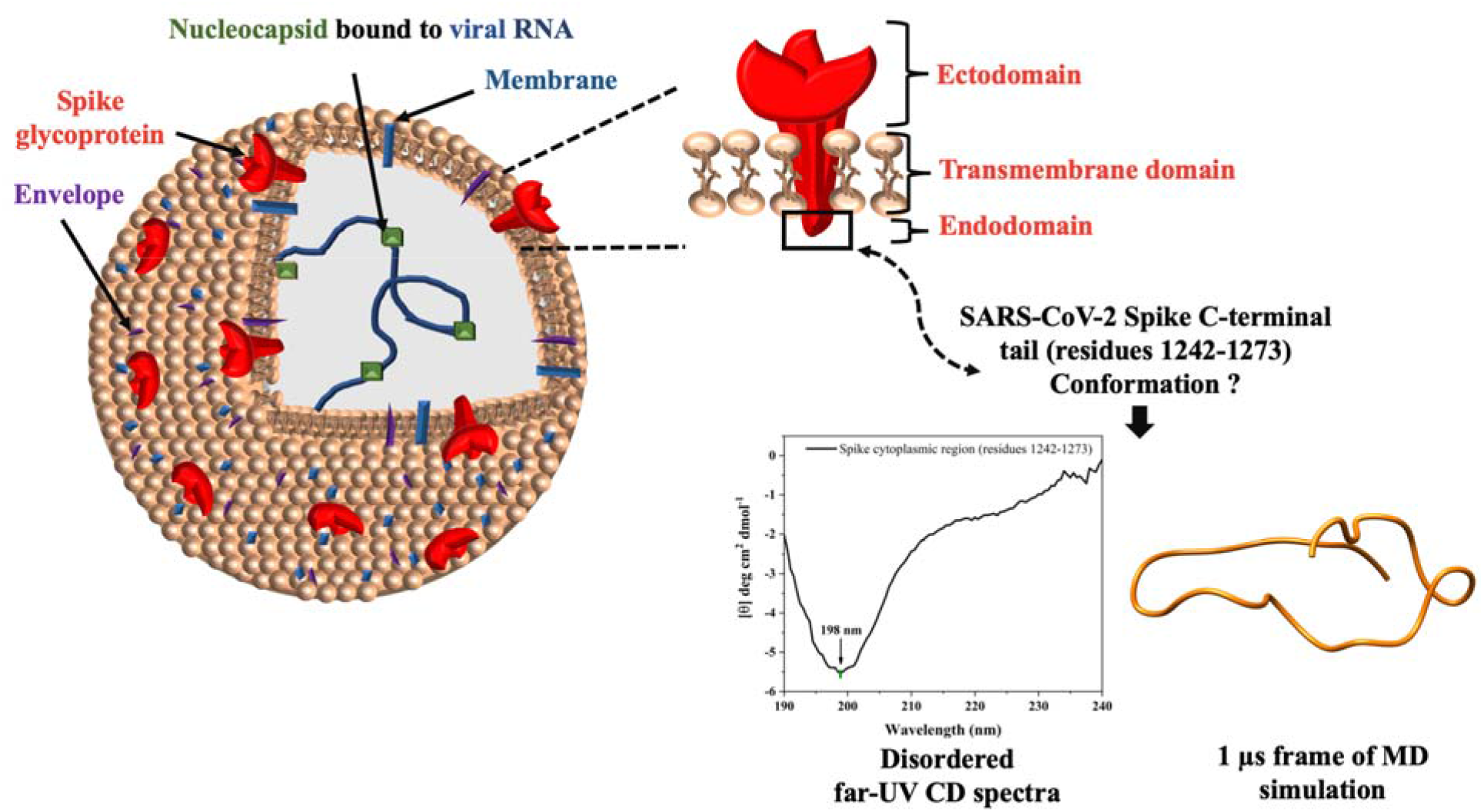

## Introduction

The importance of coronavirus spike protein is apparent from it surface-exposed location, suggesting it is a prime target after viral infection for cell-mediated and humoral immune responses as well as artificially designed vaccines and antiviral therapeutics. The SARS-CoV-2 homo-trimeric spike glycoprotein consists of an extracellular unit anchored by a transmembrane (TM) domain in viral membrane and a cytoplasmic domain [1]. It is secreted as monomeric 1273 amino acid long protein from endoplasmic reticulum (ER) shortly after which it trimerizes to facilitate the transport to the Golgi complex [1,2]. Moreover, N-linked high mannose oligosaccharide side chains that are added to spike monomer in ER are further modified in Golgi compartments [2].

Spike is one of the most extensively studied protein among all of SARS-CoV-2 proteome. So far, based on Uniprot database, approximately two hundred structures have been reported using X-ray crystallography and cryo-electron microscopy techniques. However, these structures consist of S1 subunit of spike but lacks the transmembrane and cytoplasmic C-terminal regions present in S2 subunit or with missing electron densities in cytoplasmic region. The distal S1 subunit (residues 14–685) contains a N-terminal domain, a C-terminal domain, and two subdomains (**Figure 1**). The C-terminal domain of S1 is the receptor-binding domain or RBD, has a receptor-binding motif (RBM) which interacts with human angiotensin converting enzyme 2 (ACE2), chief target receptor of SARS-CoV-2 on human cells [3]. RBM is present as an extended loop insertion which binds to bottom side of the small lobe of ACE2 receptor. The S2 subunit (residues 686-1273) has a hydrophobic fusion peptide, two heptad repeats, a transmembrane domain, and a cytoplasmic C-terminal tail (**Figure 1**).

**Figure 1:**
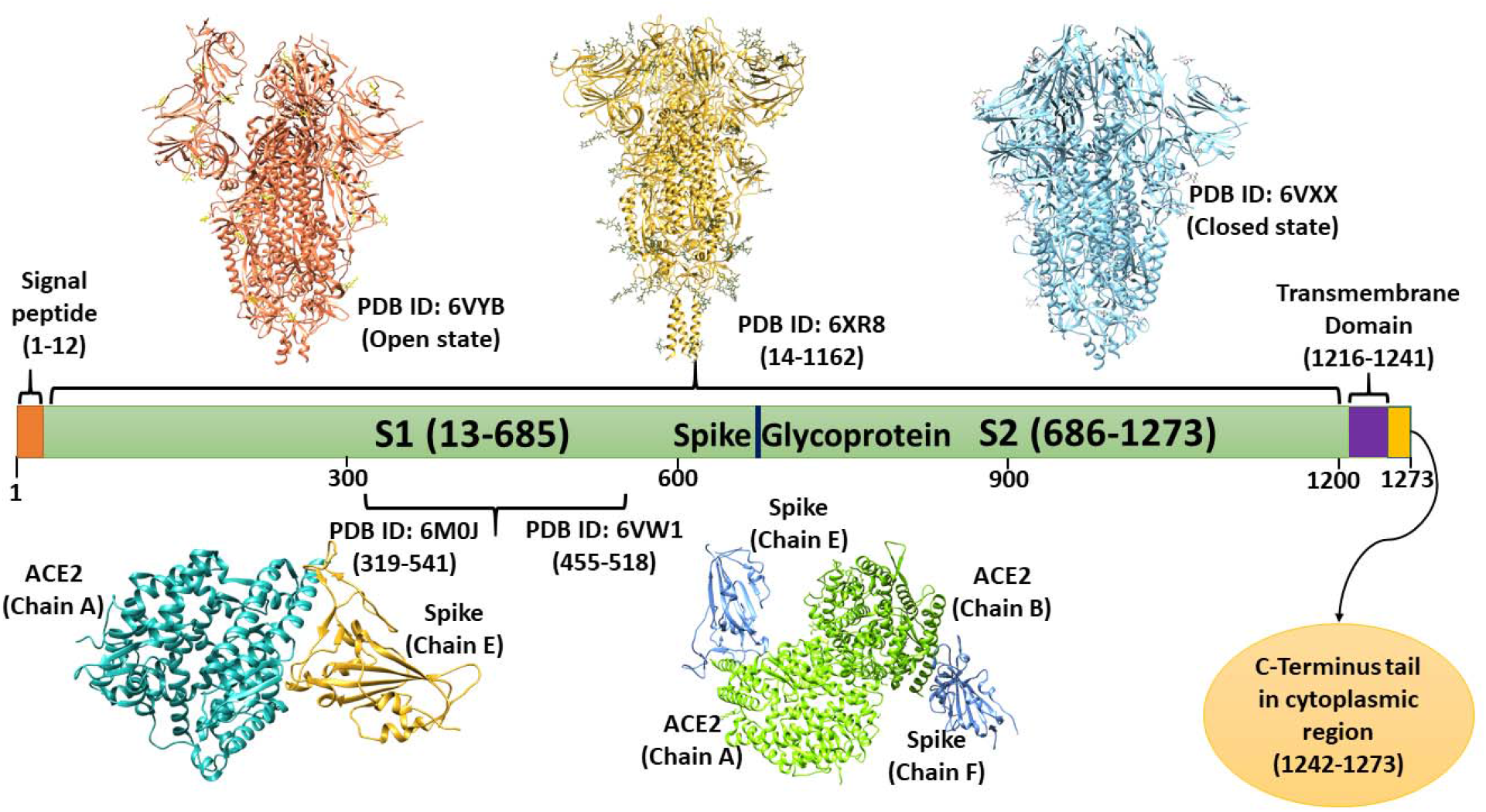
Domain architecture of spike Glycoprotein: depiction of available structures in open and closed states, transmembrane domain, and cytoplasmic C-terminal tail. Based on prediction of transmembrane region in the spike protein by CCTOP, a consensus-based predictor, the boundaries of all domains have been defined. As per CCTOP prediction, the transmembrane region of spike lies within the residues 1216-1241, and so, the cytoplasmic region of spike has been used in this study with the residues 1242-1273.

As of yet, cytoplasmic domain of spike protein is the least explored region despite of such extensive research in pandemic times. It is of particular importance as it contains a conserved ER retrieval signal (KKXX) [4]. In SARS-CoV and SARS-CoV-2 spike proteins, a novel dibasic KLHYT (KXHXX) motif present at extreme ends of the C-terminus plays a crucial role in its subcellular localization [5–7]. Also, deletions in cytoplasmic domain of coronavirus spike are implicated in viral infection in recent reports [8–11]. SARS-CoV and SARS-CoV-2 spike having a deletion of last ~20 residues displayed increased infectivity of single-cycle vesicular stomatitis virus (VSV)-S pseudotypes [8,9]. Contrarily, short truncations of cytoplasmic domain of Mouse Hepatitis Virus (MHV) spike protein (Δ12 and Δ25) had limited effect on viral infectivity while the long truncation of 35 residues interfered with both viral-host cell membrane fusion and assembly. Importantly, it is also shown to interact with the membrane protein inside host cells [10]. In our previous report, the cytoplasmic tail is predicted to be a MoRF (Molecular Recognition Feature) region (residues 1265-1273) by a predictor MoRFchibi [7]. The MoRF regions in proteins are disorder-based binding regions that contribute the binding to DNA, RNA, and other proteins. In the same report, it is also found to contain many DNA and RNA binding residues [7].

Despite of availability of several structures of spike protein using advanced techniques like cryo-EM, the structure of cytoplasmic domain is not yet clear due to its ‘missing electron density’. Generally, intrinsically disordered proteins show such characteristic of missing electron density and lacks a well-defined three-dimensional structure [12]. Additionally, the consensus-based disorder prediction by MobiDB has shown this region to be disordered [13]. Considering these arguments, we aimed to understand the cytoplasmic domain of the SARS-CoV-2 spike protein to gain further insights. To this end, we computationally analysed its behavioural dynamics using molecular dynamic (MD) simulations up to one microsecond (μs) and validated it with CD based experiments. This report’s outcomes will help understand this domain’s structure and function and provide knowledge to explore the interaction of spike protein with other viral and host proteins.

## Material and Methods

### Materials

For CD based experiments, the synthesized peptide of spike cytoplasmic region (residues 1242-1273) with purity 85.3 % was procured in lyophilized form from GeneScript, USA. Chemicals and solvents i.e., 2,2,2-Trifluoroethanol (TFE) and Sodium Dodecyl Sulfate (SDS) with more than 99% of purity were purchased from Sigma-Aldrich, USA. In addition, macromolecular crowding agents, polyethylene glycol (PEG 8000), and sucrose are used to examine the structural behavior of spike-CTR in cell-like crowding conditions. Further, the lyophilized peptide of spike cytoplasmic region was dissolved in nuclease free water with a concentration of 1 mg/ml and prepared a stock concentration of 289.35 μM for all experimental measurements.

#### Transmembrane prediction

Before studying the cytoplasmic domain, we have applied multiple servers to predict the spike protein’s transmembrane region. TMHMM [14], TMPred [15], SPLIT [16], PSIPRED [17,18], and CCTOP [19] web predictors works using highly optimized and least biased algorithms which also takes into account the homology of sequences. Among the predictors, CCTOP predicts the transmembrane regions based on the consensus of multiple predictors and experimental derived structural information from homologous proteins in the database. Therefore, it provides a better understanding and identification of transmembrane regions in the protein. Based on CCTOP prediction, we have chosen the C-terminal cytoplasmic tail region to elucidate its structural dynamics.

#### Disorder Prediction

The propensity of disorderedness in spike C-terminal cytoplasmic tail region is predicted using PONDR family [20–22], IUPred long [23], and PrDOS [24] servers. The detailed methodology is given in our previous reports [7,25].

#### Structure Modelling

PEP-FOLD 3.5 webserver [26] is used to predict 3D structures of selected spike protein regions. By implementing *optimized potential for efficient structure prediction* (OPEP) coarse-grained forcefield-based simulations, an improved and minimized structure is obtained as described earlier [27,28]. Then, the structure is prepared in Schrodinger suite where the missing hydrogens, improper bond orders, and protonation states are corrected. Further, the prepared structure is used for MD simulations.

#### Molecular Dynamic (MD) Simulations

To comprehend and comparable outcomes for intrinsically disordered regions, we have used two different forcefields to analyze the structural dynamics of the cytosolic domain of spike protein.

##### Simulation with OPLS 2005

We have used Desmond simulations package, where simulation setup is built by placing the protein structure in an orthorhombic box along with TIP3P water model, 0.15M NaCl salt concentration [29]. After solvation, the system is charge neutralized with counterions using OPLS 2005 forcefield. To attain an energy minimized simulation system, the steepest descent method is used for 5000 iterations. Further, the equilibration of system is done to optimize solvent in the environment. Using NVT and NPT ensembles within periodic boundary conditions, the system is equilibrated for 100 ps each. The average temperature at 300K and pressure at 1 bar are maintained using Nose-Hoover thermostat and Martyna-Tobias-Klein (MTK) coupling methods during simulation [30,31]. All bond-related constraints are solved using SHAKE algorithm, and hydrogen bond constraints are solved using LINCS algorithm [32]. The final production run is performed for 1 μs using our in-house facilities.

##### Simulation with CHARMM36m

Another forcefield we used, CHARMM36m in Gromacs, is an improved version of CHARMM36, which is effectively developed for analyzing IDP regions in the proteins in significant simulation timescale [33,34]. Using TIP3P water model, the system is prepared for proper electrostatic distribution and then neutralized for charge using counterion (1 Na+ ion in this case). The energy minimization of simulation setup using steepest descent method is done for 50,000 steps. For temperature and pressure coupling, the V-rescale and Parrinello-Rahman algorithms are used where 300K and 1 bar is the average temperature and pressure respectively. After equilibration of 100 ps for NVT and NPT methods, the production run is then executed for 1 μs using our high performing cluster at IIT Mandi.

##### Replica-Exchange Molecular Dynamic (REMD) simulations

The enhanced conformation sampling using REMD simulations is widespread in protein folding. During REMD simulations, the swapping of conformations occurs and reduces the chances of entrapping simulations in local minimum energy states [35]. Therefore, we have performed REMD using eight replicas (numbered from 0 to 7) at temperatures 298 K, 314 K, 330 K, 346 K, 362 K, 378 K, 394 K, and 410 K, calculated by linear mode of Desmond. The last frame of 1 μs of Desmond simulation trajectory is chosen as the initial conformation for REMD. The multigrator integrator of Langevin and Nose-Hoover as thermostats, whereas Langevin and Martyna-Tobias-Klein (MTK) are used to equilibrate the systems [30,31]. The accountability of conformation swaps to be accepted or rejected is done based on common Metropolis criterion using the following equation:

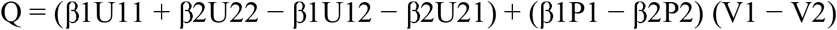

Where *U_ij_* = potential energy of replica *i* in the Hamiltonian of replica *j*,

*P_i_* = the reference pressure of replica *i*,

*V_i_* = instantaneous volume of replica *i*, and

*β_i_* = the inverse reference temperature of replica *i*.

If Q > 0 = accept, or Q < −20 = reject the exchange,

else accept the exchange if *rand_N_* < exp(Q), where *rand_N_* is a random variable on (0,1).

#### CD spectroscopy

To obtain the secondary structure based spectral information, we employed JASCO CD instrument (Jasco J-1500 CD spectrometer, USA). 25 μM of spike cytoplasmic region was prepared in 20 mM sodium phosphate buffer (physiological pH 7.4) to record the spectra in isolation. Further, to observe the structural changes in peptide, CD spectra (with same concentration) was recorded in increasing concentration of organic solvent TFE (0 to 90%). Similarly, conformational changes in peptide were monitored in presence of an anionic detergent, Sodium Dodecyl Sulfate (SDS) (below and above critical micellar concentration (CMC)). Additionally, the intracellular crowding environment mimicking agents PEG 8000 and Sucrose were utilized (at concentrations from 50-300 g/L) to examine the structural changes in the peptide. All spectra were recorded in far-UV region from 190 nm–240 nm in 1 mm quartz cuvette. The scan speed was kept at 50 nm/min with a response time of 1s and 1 nm bandwidth. With similar parameters, baseline spectra of buffers were recorded and subtracted from all spectral measurements. In case of macromolecular crowders experiments, the baseline spectra consisted buffers and crowding agents. The resultant data were plotted as wavelengths vs ellipticity (with HT value, < 600 volts) at x and y axis respectively using Origin software.

## Results

In recent times, computational approaches have been widely used to explore the secondary and tertiary structures of proteins and small peptides. It has immensely helped in the ongoing COVID-19 pandemic to study structural conformations of protein and their interacting partners, i.e., protein, ligands, glycans, etc. Molecular dynamics simulation is a useful approach to answer such questions at the atomic level. Herein, we have studied the cytoplasmic domain and performed rigorous simulations to unravel the structural dynamics of least explored cytoplasmic domain of essential spike protein.

### Transmembrane region analysis

The sequence-based analysis of transmembrane region and disorder prone regions have also been analyzed. The subcellular localization of spike protein occurs in the extracellular, transmembrane, and cytoplasmic regions [36]. However, based on SARS-CoV and SARS-CoV-2 proteins sequence alignment, approximately 77% similarity is found among both viruses spike proteins [7]. The C-terminal has shown high similarity and conserved regions, while the N-terminal has vastly varying residues.

Based on multiple predictors used in this study, spike protein’s transmembrane region lies within 1213-1246 residues (**Figure 2**). A consensus-based server, CCTOP, has predicted the transmembrane region from residues 1216-1241, which is more reliable as it compares and uses the previously available experimental information of related proteins. Therefore, the cytoplasmic region is selected from 1242 to 1273 amino acids (sequence: NH2-SCLKGCCSCGSCCKFDEDDSEPVLKGVKLHYT-COOH).

**Figure 2:**
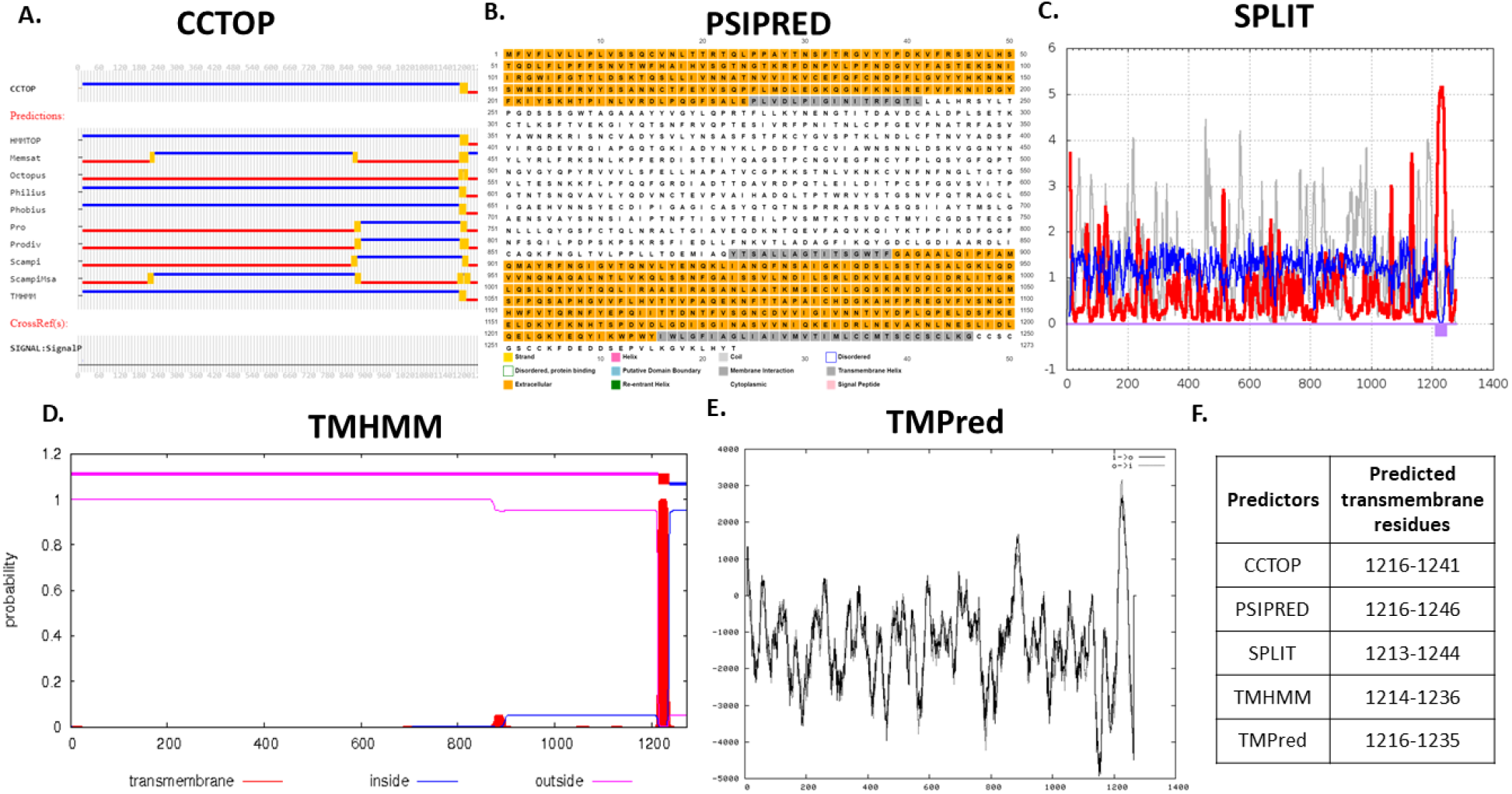
Transmembrane region prediction from five web servers: **A.** CCTOP, **B.** PSIPRED, **C.** SPLIT, **D.** TMHMM, and **E.** TMPred. **F.** Table showing predicted transmembrane residues. All predictors work as standalone except CCTOP which works based on consensus form multiple predictors. Considering this, the cytoplasmic region of spike is chosen from residues 1242-1273 as CCTOP has predicted residues 1216-1241 in transmembrane region.

### Disorder prediction

In our recent study, we have identified the disordered and disorder-based binding regions in SARS-CoV-2 where the cytoplasmic domain at C-terminal of spike protein is found to be disordered [7]. Again, we analyzed the disorderedness in selected cytoplasmic region using multiple predictors, including PONDR family, IUPred long, and PrDOS predictors. Out of six predictors, three predictors from PONDR family have predicted it as highly disordered, PrDOS has predicted it as moderately disordered. In contrast, PONDR FIT and IUPred long predicted it as least disordered (**Figure 3**).

**Figure 3:**
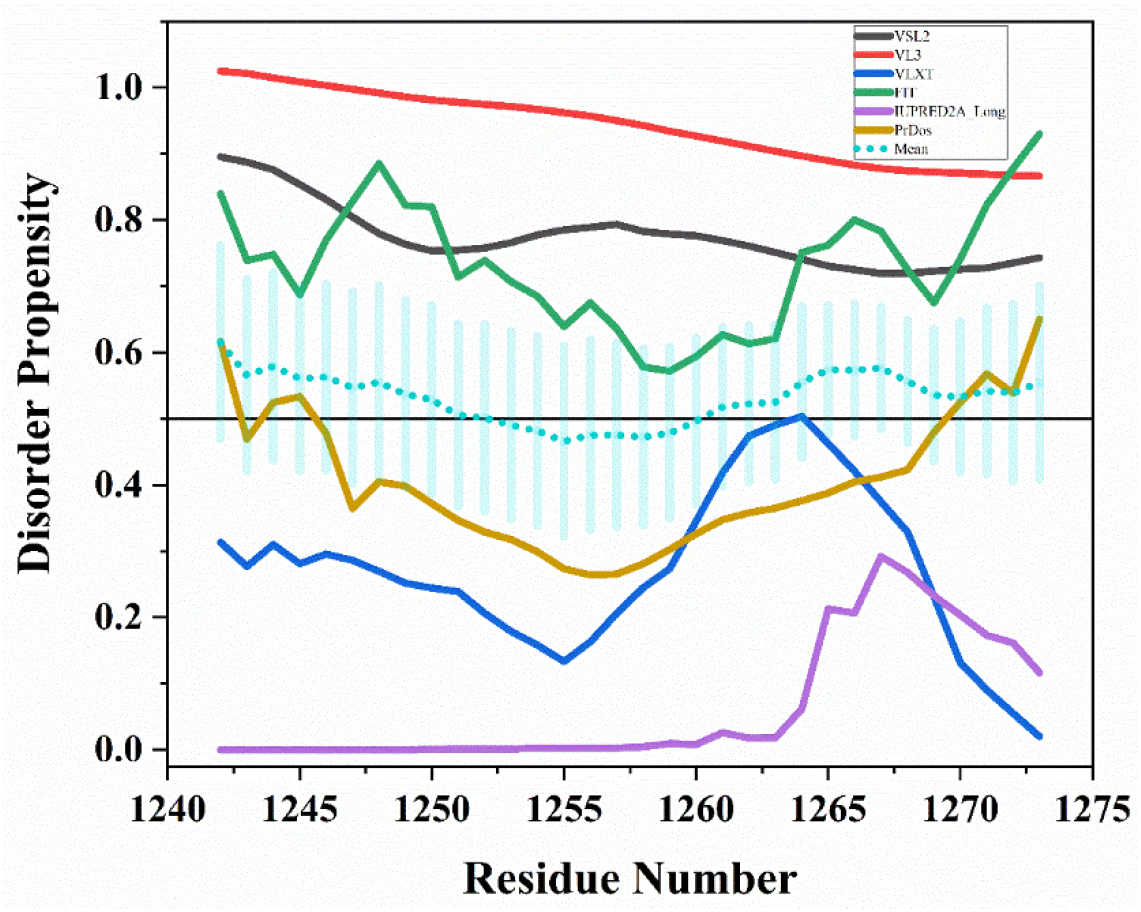
Intrinsic disorder analysis of spike C-terminal cytoplasmic tail (residues 1242-1273) region using six predictors including PONDR family (VSL2, VL3, VLXT, and FIT), IUPred long, and PrDOS servers. The mean line is denoted in short dots style, and the standard error bars on mean are also highlighted.

### Structure modelling with PEPFOLD3

In absence of an experimentally determined 3D structure of protein, structure modelling provides an approximate structure model based on homology and properties of amino acids using a wide range of optimized algorithms. It generates the prototype fragments and assembles them by implementing *optimized potential for efficient structure prediction* (OPEP) coarse-grained forcefield-based simulations. The best-obtained structure of extreme C-terminus tail (1242-1273 amino acids) containing two small helical regions is further prepared for MD simulations in aqueous conditions. These helical regions are present at residues _**1243**_CLKGC_**1247**_ and _**1265**_LKGV_**1268**_ of spike glycoproteins C-terminus (**Figure 4A**).

**Figure 4:**
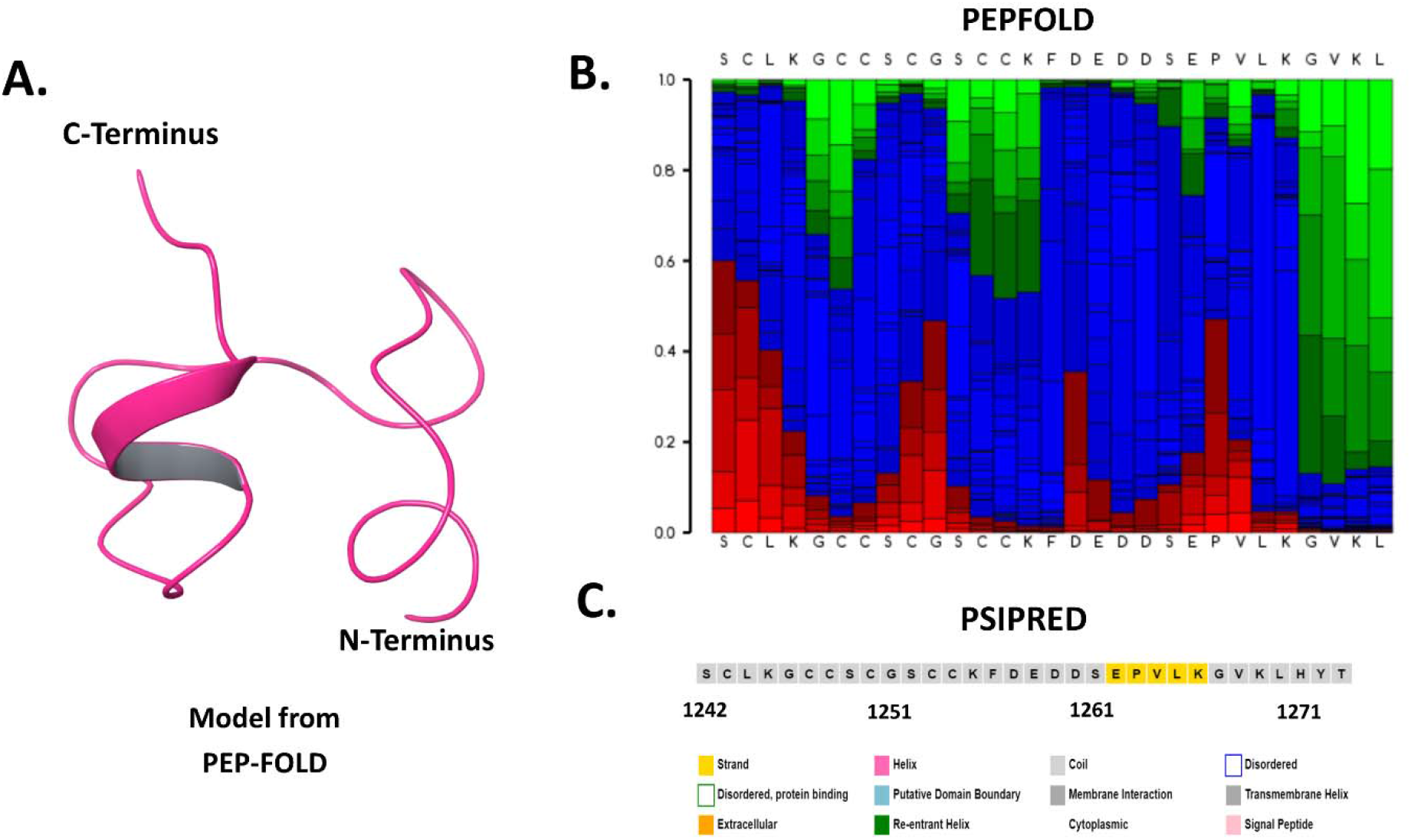
Sequence and fold propensity analysis of spike C-terminal cytoplasmic domain (1242-1273 residues): **A.** Modelled structure through PEP-FOLD web server, **B.** PEP-FOLD model structure analysis depicting helix (red), coil (blue), and extended (green), and **C.** Secondary structure analysis using PSIPRED web server.

### Simulation with OPLS 2005

In the last three decades, many advancements have been made in forcefields and hardware related to MD simulation to match the experimental events. Long MD simulations up to microseconds or milliseconds are incredibly insightful to study structural conformations occurring at the nanoscale level. We have recently explored various regions of different SARS-CoV-2 proteins through computational simulations and experimental techniques that are very well correlated [28,37]. This study performed 1 μs MD simulations of C-terminal cytoplasmic domain of spike protein (1242-1273 residues) to understand its dynamic nature. As obtained from structure modelling through PEP-FOLD, the model contains two small helices at both terminals with residues _**1243**_CLKGC_**1247**_ and _**1265**_LKGV_**1268**_ (**Figure 4A & 4B**). According to *2struc* webserver [38], these helices contribute to 12.5% of total secondary structure while rest of the region is constituted by turns and extended coils. The secondary structure prediction of spike C-terminal tail region contains a -strand of five residues _**1262**_EPVLK_**1266**_, as predicted by PSIPRED webserver [17] (**Figure 4C**).

After analyzing the disorder propensity and secondary structure composition, we performed a rigorous simulation of cytoplasmic region (residues 1242-1273) to understand its atomic movement and structural integrity. A total of 1 μs simulation was done after 50,000 steps of steepest descent method-based energy minimization. It has been observed that the structure of spike C-terminal cytoplasmic region remains to be unstructured throughout the simulation. Based on mean distance analysis, the peptide simulation setup showed massive deviations up to 7.5 Å which does not attain any stable state (**Figure 5A**). As shown in **figure 5B**, mean fluctuation in residues is observed to be within the range of 1.6 – 6.4 Å. The intramolecular hydrogen bond analysis demonstrates the highly fluctuating trend portraying no stable helical or beta sheet conformation adoption by the residues (**Figure 5E**). The secondary structure timeline (**Figure 5C & 5D**) also reveals the disordered state of spike C-terminal cytoplasmic region during the 1 μs simulation time (none of the frames captured α-helix or β-sheets) which is further depicted in the snapshot of 1 μs frame in **figure 5F** and the trajectory movie upto 1 μs (**supplementary movie 1**).

**Figure 5:**
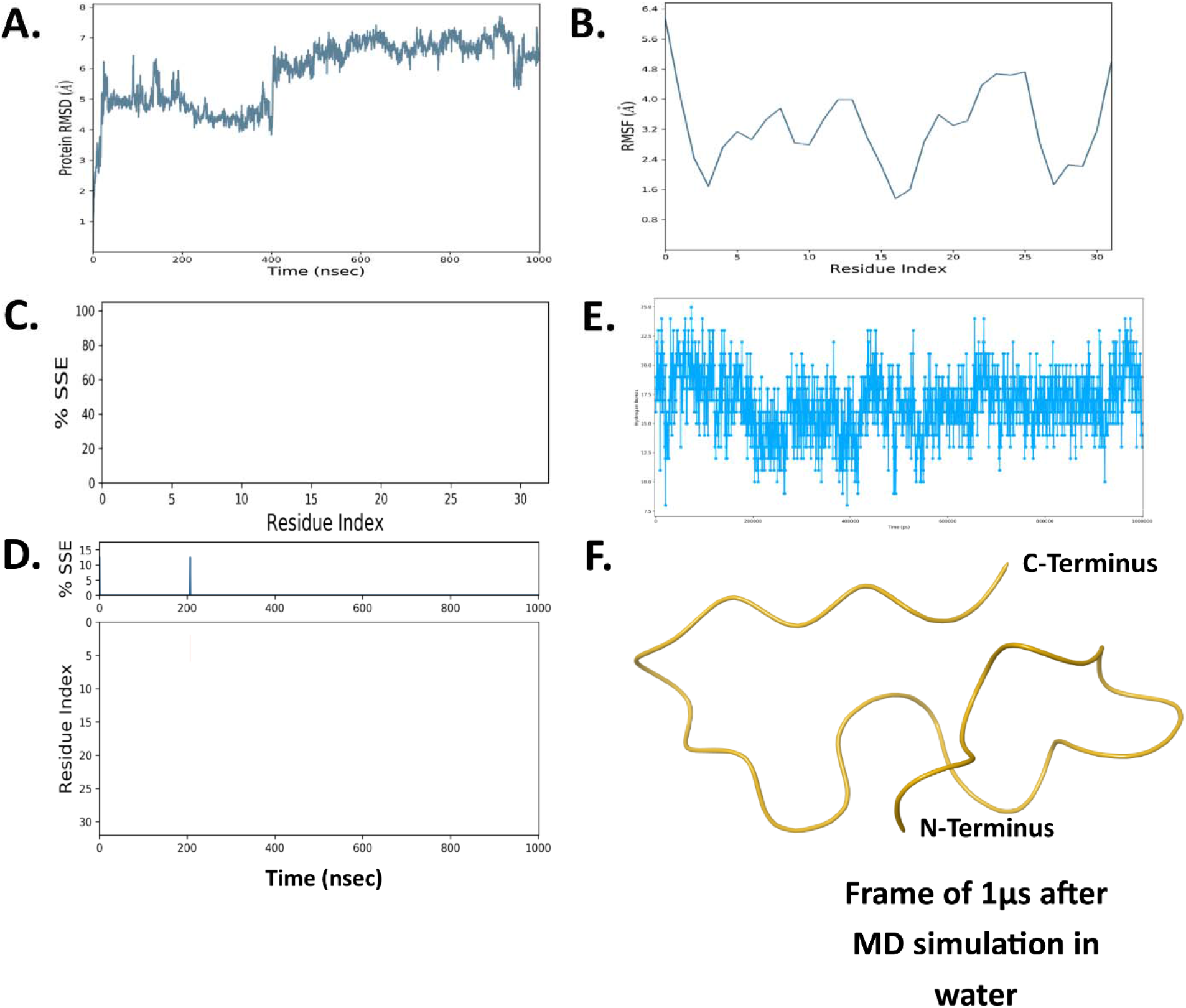
One microsecond MD Simulation analysis of spike C-terminal cytoplasmic domain (1242-1273) using OPLS 2005 forcefield: **A.** Root mean square deviation (RMSD), **B.** Root mean square fluctuation (RMSF), **C.** Secondary structure element (SSE) of residues, **D.** Timeline representation of secondary structure content during 1μs simulation time, **E.** Hydrogen bonds of protein and **F.** Last frame at 1 μs.

We have also modelled the cytosolic part of Spike protein from 1235-1273 residues as defined in Uniprot database and two predictors (TMPred: 1216-1235, and TMHMM: 1214-1236) used in this study. In modelled structure, the helical propensity in cytosolic region was shown by 1237-1245 residues. Using above described OPLS 2005 forcefield parameters, the all-atoms explicit solvent MD simulation was carried out for 1 μs. The trajectory analysis has been shown in **supplementary figure 1**, the cytosolic region has revealed majorly unstructured region along with a small β-strand of two residues _1258_FD_1259_ after 1 μs. The upward trend of RMSD values illustrates the highly deviating atomic positions and fluctuating RMSF shows the change in structural property of residues (**supplementary figure 1A & 1B**). Also, the decreasing number of hydrogen bonds demonstrates the breaking of helices in the structure (**supplementary figure 1C**). The time-dependent secondary structure element analysis illustrates that a total of 15% secondary structure was formed that includes mainly alpha helix and small percentage of beta strands (**supplementary figure 1D & 1E**; red: alpha helix and blue: beta strands). After huge structural transitions, the structural composition of last frame of simulation is shown with a small beta strand of two residues and other regions to be disordered (**supplementary figure 1F**). The snapshots at every 100 ns till 1 μs show the structural transitions in Spike cytoplasmic region (**supplementary figure 2**).

### Desmond trajectory clustering

In a microsecond simulation trajectory, a large amount of data can be sampled and simply understood using techniques like clustering [39]. Desmond applies affinity propagation algorithm to produce the representatives of each identified cluster [40]. It calculates a similarity matrix for sampling the frames of trajectory. Here, we have done trajectory clustering based on RMSD values of backbone atoms of Spike cytoplasmic tail region with a timer threshold of 0.2. A total 15 clusters with size 39 to 4 structures were identified, out of which top 10 are shown here in **figure 6**. The RMSD values of these identified structures in reference to the first frame of trajectory is in the range of 4.53 to 6.85 Å. All cluster representative structures are fully unstructured and show no propensity for helical or beta sheet conformations in aqueous condition.

**Figure 6:**
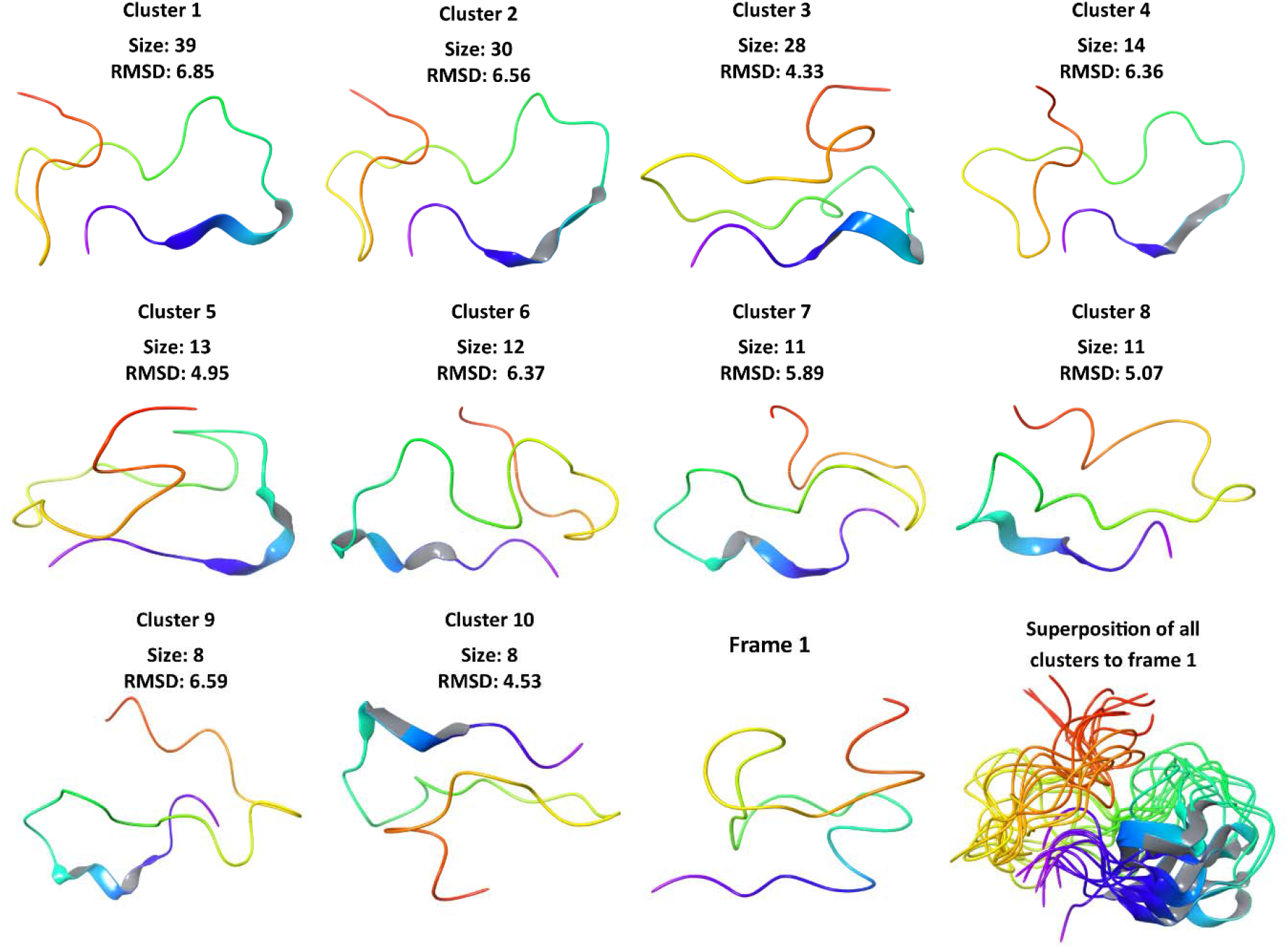
Depiction of representative from top 10 clusters from 1 μs simulation trajectory. Size of each cluster is shown which represent the total number of frames in the trajectory based on RMSD calculated with reference structure (frame 1). The protein backbone of all frames is shown as superimposition.

### Simulation with CHARMM36m

The characterization of intrinsically disordered proteins (IDPs) and regions (IDPRs) using MD simulations is now very well persistent in literature. In last two decades, several forcefields for MD simulations are developed and improved at times. All forcefields have their advantages and limitations due to the proper evaluation of secondary structure composition. Here, we have used another forcefield, CHARMM36m, for determining the conformational dynamics of spike cytosolic region. As shown through trajectory snapshots at every 100 ns in **figure 7**, the cytosolic region has adopted a β-sheet conformation at its N-terminal. Two β-strands can be seen with varying amino acid length at every 0.1 μs frame. The only exceptions are 0.3 μs and 0.4 μs frames, which do not show any secondary structure. Further, after 0.5 μs frame, a gradual loss in two β-strands indicates a gain in disorder content in spike C-terminus. To this end, the 1 μs simulation frame comprises of only a short β-strand at residues _1257_KFD_1259_ and rest unstructured residues.

**Figure 7:**
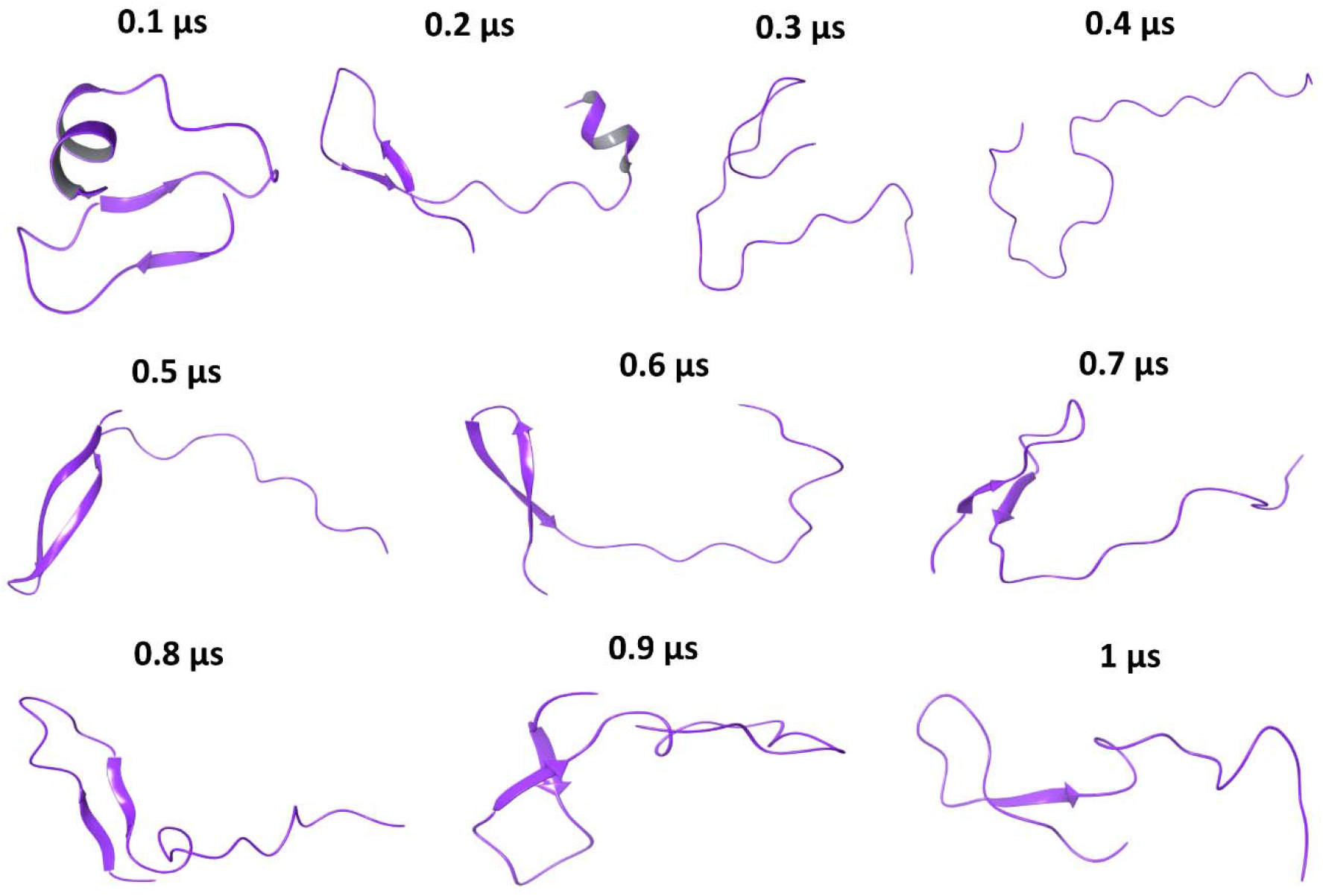
MD simulation with CHARMM36m forcefield: Snapshots at every 100 ns of 1 μs simulation trajectory.

These structural changes have been the reason for immensely varying atomic distances throughout the simulation. Likewise, RMSD values are found in a range of approx. 0.75 nm to 1.75 nm (**Figure 8A**); the mean residual fluctuations are in the range of 0.5 nm to 1.1 nm (**Figure 8C**), and the Rg values vary up to 2.4 nm (**Figure 8B**), which demonstrates the vastly changing structural compactness. The timeline representation of secondary structure composition at each frame also depicts the structural inconsistency throughout the simulation (**Figure 8E**). As calculated in VMD [41], the total number of salt bridges are found to be 11 in the trajectory between the residues Asp1257-Lys1269, Glu1258-Lys1266, Glu1258-Lys1245, Asp1259-Lys1269, Glu1262-His1271, Glu1262-Lys1266, Glu1262-Lys1269, Asp1260-Lys1266, Asp1259-His1271, Asp1259-Lys1266, and Asp1260-Lys1269. The free energy landscape of complexation of Spike cytosolic region with variables RMSD and Radius of gyration is calculated using a Python script *generateFES.py* (http://www.strodel.info/index_files/lecture/html/media/generateFES.py) which reveals that the most of the conformations are lying with least energy from 0 to 4 kcal/mol. Convincingly, it is evident that a major part of spike cytosolic region is disordered.

**Figure 8:**
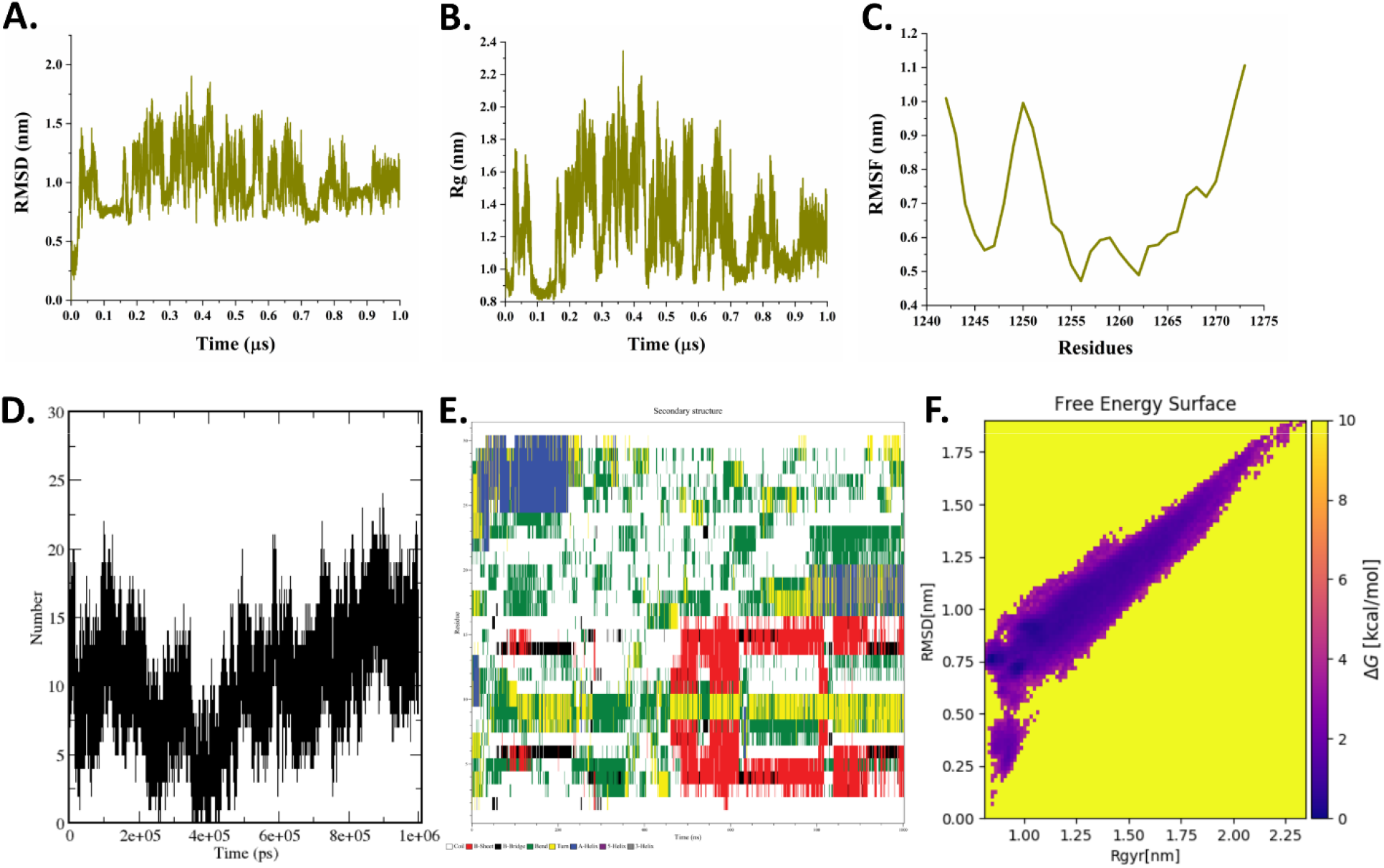
Trajectory analysis of spike cytoplasmic region with CHARMM36m forcefield: **A.** RMSD, **B.** Radius of gyration (Rg), **C.** RMSF, **D.** Hydrogen bonds and **E.** Secondary structure element timeline during the course of 1 μs simulation time, and **F.** Free energy landscape of simulation trajectory using RMSD and Radius of gyration as variables.

### Conformation sampling using Replica-Exchange Molecular Dynamic (REMD) simulations

The disordered form (last frame) of spike cytoplasmic region from Desmond simulation trajectory is used for REMD at 8 temperatures *viz*. 298 K, 314 K, 330 K, 346 K, 362 K, 378 K, 394 K, and 410 K upto half a microsecond using 8 replicas (numbered as 0 to 7). As shown here in **figure 9**, the cytoplasmic region has adopted a β-sheet structure at increasing temperatures upto 394 K. In comparison, at 410 K in replica 7 these changes are found to be reversible. Although previous frames of replica 7 display the formation of multiple long β-strands throughout simulation time. According to snapshots illustrated in **figure 9**, the cytosolic region has gained three to four β-strands and appears to be in a well-folded manner as temperature increases.

**Figure 9:**
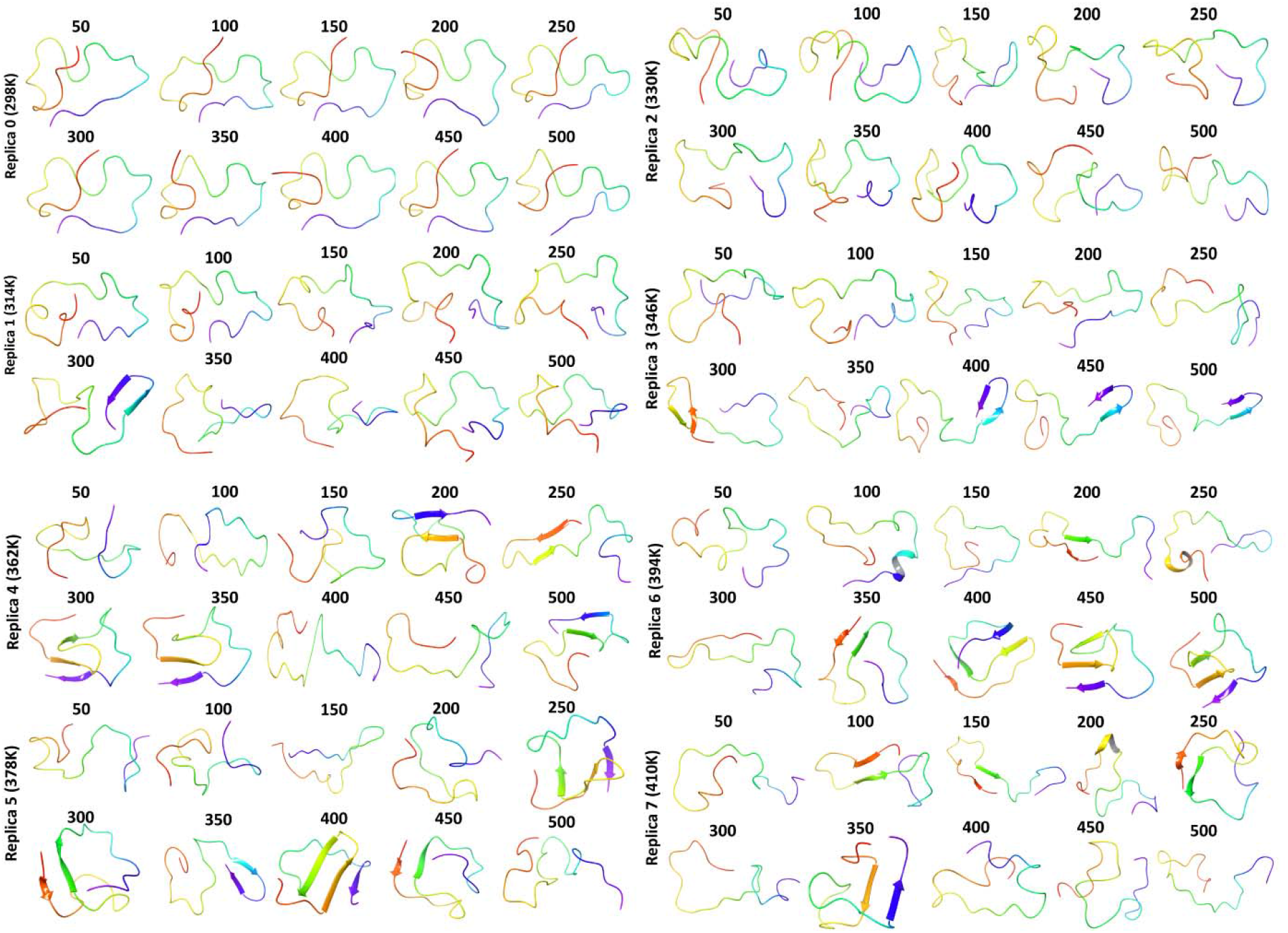
Snapshots from all replicas at every 50 ns during half a microsecond REMD simulation.

For a clear understanding of all frames in REMD, timelines of all 8 replicas are displayed in **figure 10** where the consistent formation of β-strands due to rising temperature is also validated. However, at extreme temperature (410K), some structural changes are reversed, and the total secondary structure element (SSE) gets reduced in comparison to previous replicas (**Figure 10 Table**). According to mean distances analyses, huge fluctuation is observed in replicas 3, 5, and 6 where structural changes occurred (**Figure 11A, 11B, 11D**). As elucidated through hydrogen bond analysis (**Figure 11C**), the highest numbers are 21, 20, 20 for replicas 2, 6, and 7, respectively. The superimposed last frame of each replica shows structural differences with atomic distances (RMSD) in range of 3.7 Å to 8.5 Å with respect to the starting frame for REMD (**Figure 11E**).

**Figure 10:**
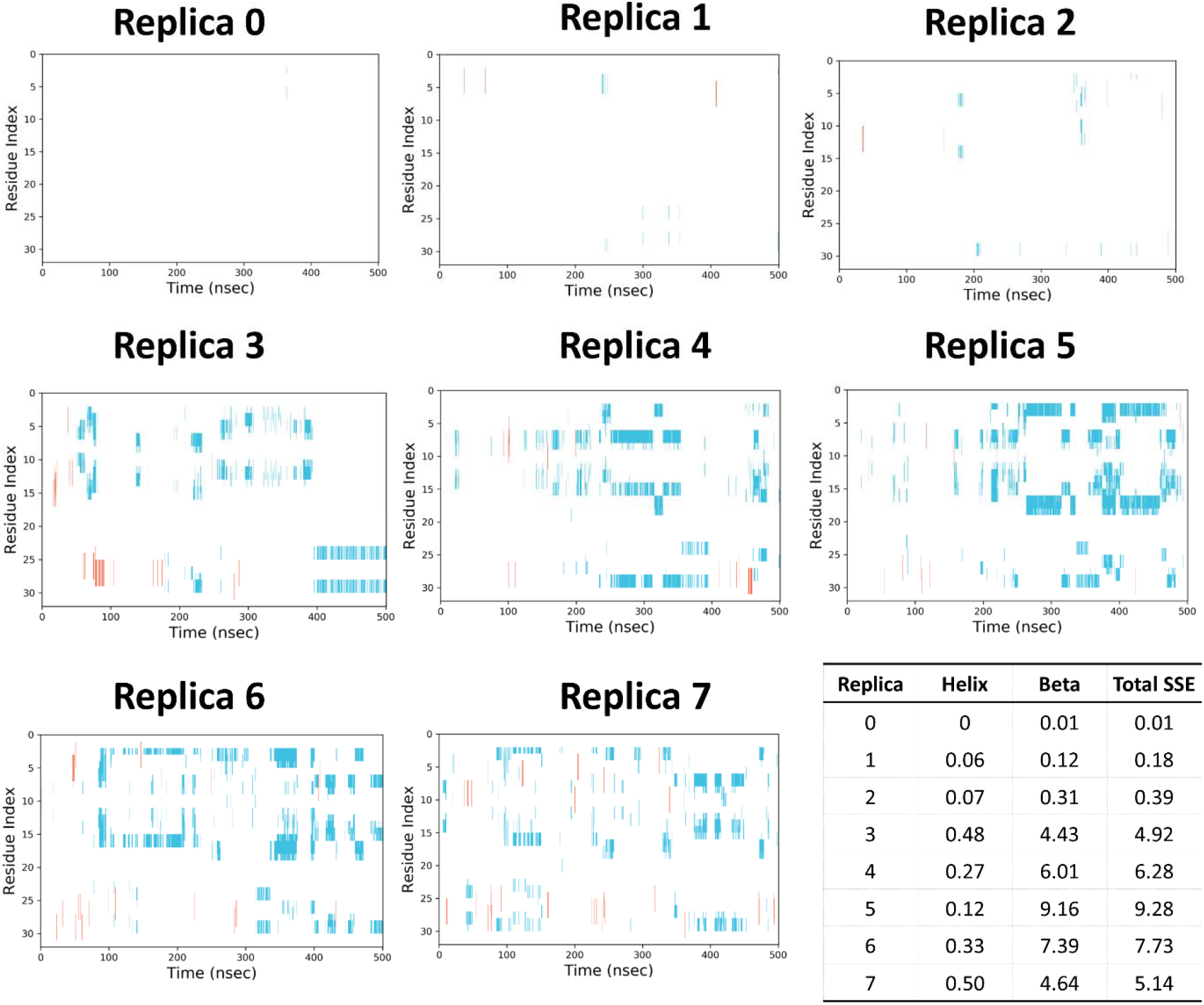
Timeline representation of simulation trajectories from all replicas upto half a microsecond simulation time. A table of percentage secondary structure elements in each simulation.

**Figure 11:**
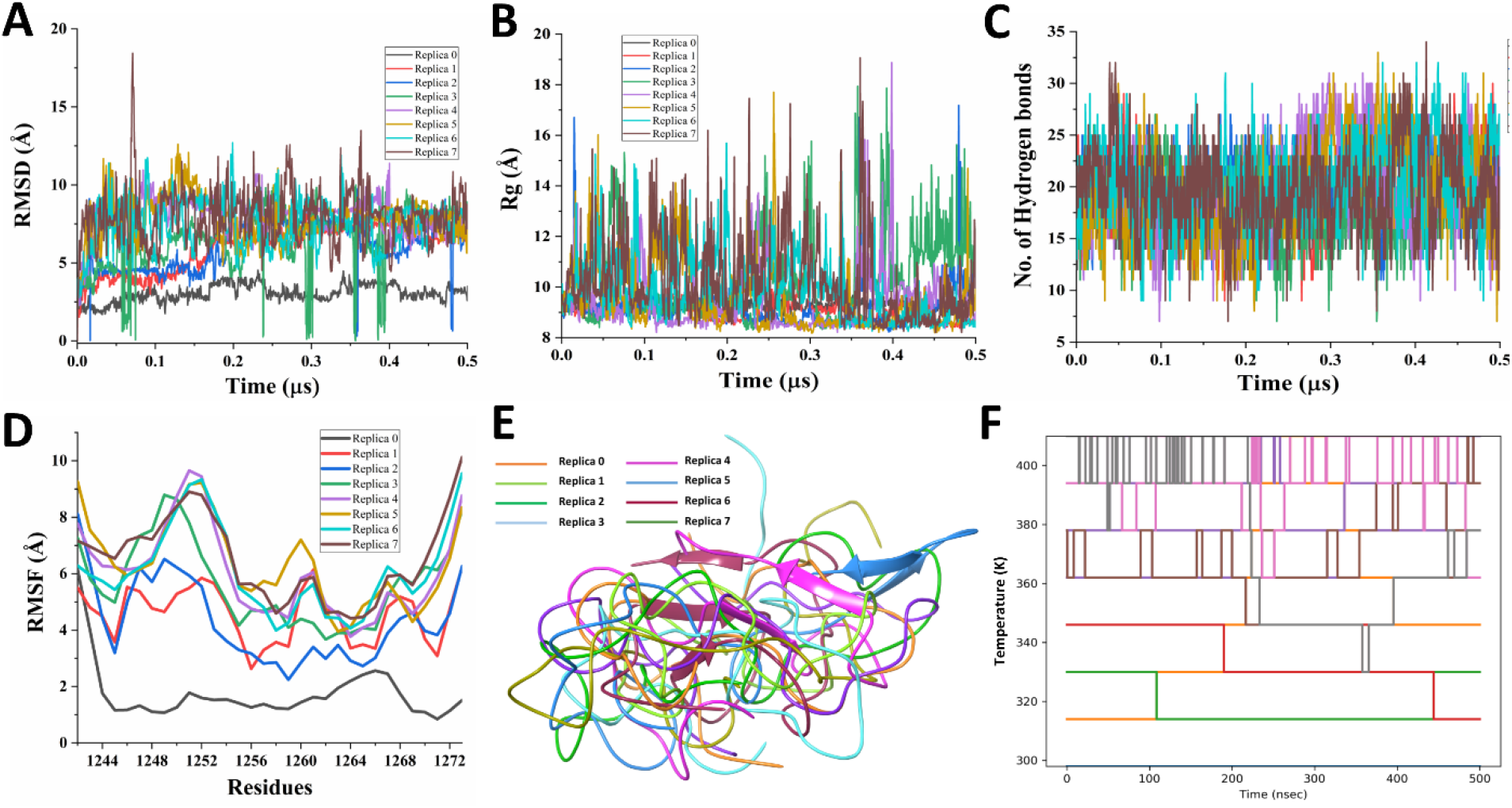
Trajectory analysis from all replicas simulated at different temperatures: **A.** RMSD, **B.** Rg, **C.** Number of hydrogen bonds, **D.** RMSF, **E.** Superimposed last frames, and **F.** Conformation exchange review during REMD.

## CD spectroscopy analysis

### Spike cytoplasmic region is unstructured in isolation

As observed through MD simulations using two different forcefields, the spike cytoplasmic region (residues 1242-1273) is unstructured or disordered. To validate our findings experimentally, we have performed CD spectroscopy-based experiments of the synthesized peptide of same region at 25 μM concentration in sodium phosphate buffer at physiological pH 7.4. In a good correlation with computer simulations, the spike cytoplasmic region has been confirmed to be disordered with a signature negative ellipticity peak at 198 nm (**Figure 12**).

**Figure 12:**
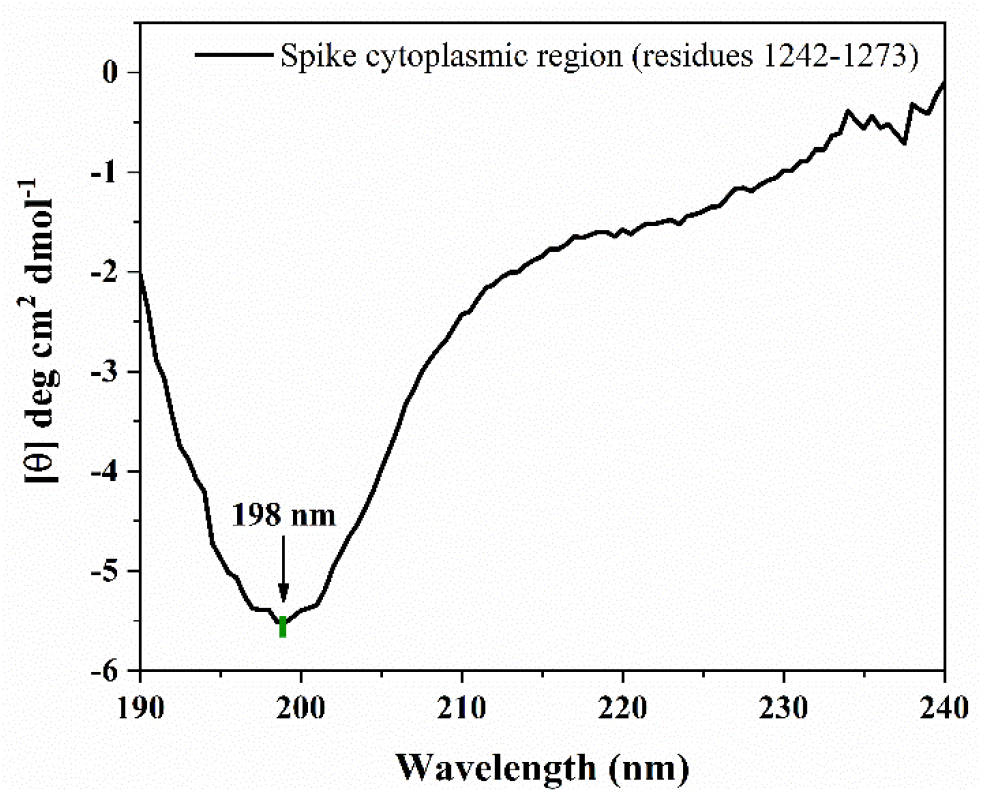
Disordered nature of spike cytoplasmic region (residues 1242-1273): Far UV-CD spectra of 25 μM spike cytoplasmic region in presence of 20mM sodium phosphate buffer (pH 7.4) is recorded from 190 nm to 240 nm. The significant negative ellipticity at 198 nm is characteristic of a disordered conformation.

Further, we have also investigated the gain-of-structure property of spike cytoplasmic region. For this purpose, we have employed a well-known secondary structure inducer, 2,2,2-trifluoroethanol (TFE). TFE weakens H-bonds between NH and CO of the protein backbone with surrounding H_2_O and if observed to stabilize the intra-chain H-bonds during secondary structure formation [42]. In presence of the organic solvent TFE, spike cytoplasmic region (at 25 μM) is observed to attain an α-helical structure. The peptide slowly started bending towards the hallmark helical negative peaks at 208 nm and 222 nm before 30% of TFE concentration (**Figure 13A** and **13B**). After increment to 40% TFE, the helical peaks having negative ellipticity at 208 nm and 222 nm are significantly clear representing the structural transformation of spike terminal tail from disordered to α-helical indicating its gain-of-structure property. Although at this moment we have no clues if such things happen in real biological scenario.

**Figure 13:**
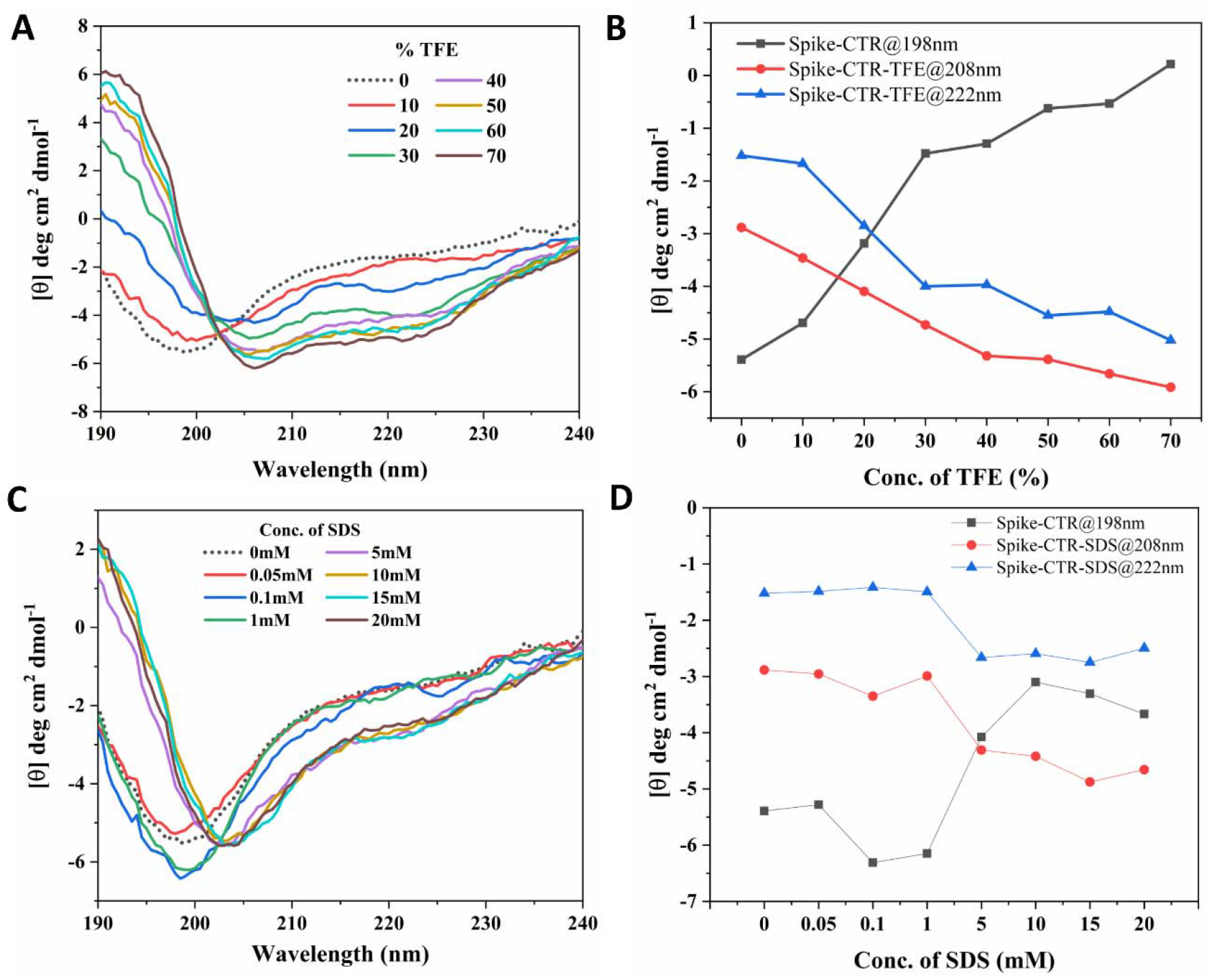
Secondary structure analysis of spike cytoplasmic region (residues 1242-1273) using CD spectroscopy: **A.** Gain in secondary structure of spike cytoplasmic region in presence of different concentrations of a secondary structure inducer TFE (2,2,2-trifluroethanol). The spike cytoplasmic region started gaining helical structure (negative ellipticity peaks near to 208 nm and 222 nm) around 30% of TFE concentration. **B.** Plot showing the change in ellipticity at 198 nm, 208 nm and 222 nm in CD spectra of spike cytoplasmic region in presence of TFE **C.** In presence of SDS (Sodium Dodecyl Sulfate), no prominent change in unstructured nature is noticed. **D.** Plot represents the change in ellipticity at 198 nm, 208 nm and 222 nm in CD spectra of spike cytoplasmic region in presence of SDS.

Next, we used sodium dodecyl sulfate (SDS), an artificial membrane-mimicking micelle forming ionic detergent to evaluate the fundamental gain-of-structure property of spike C-terminal tail. It is known to bind positively charged and hydrophobic residues of proteins using its alkyl chains and sulphate groups [43]. Contrarily to TFE, on addition of SDS, no significant change in disordered nature is observed. As evident from **Figure 13C and 13D**. not much gain in negative ellipticity at 208 nm and 222 nm is detected using CD spectroscopy.

### Spike cytoplasmic region shows no change in presence of macromolecular crowders

It is a known fact that intracellular region is highly crowded and occupies a volume of 5-40 % in the cell [44]. Crowding conditions can substantially affect several thermodynamic processes such as protein folding, change of conformation, and protein aggregation, etc. [44]. Therefore, in order to extrapolate the cellular conditions and behaviour of spike cytoplasmic tail in crowded environment, we have used two widely employed macromolecular crowders PEG 8000 and sucrose to investigate the change in conformation of spike endodomain. Here, according to our observations, in presence of either of the crowders at high concentrations up to 300 g/L (generally 80-400 g/L in cells [45]), no change in secondary structure is observed. At all concentrations of PEG 8000, the negative ellipticity at 198 nm is seen to be similar to disordered spectra with only a minimal shift in wavelength. Whereas, in presence of another crowder, sucrose, an increment in negative ellipticity from −2 deg cm^2^ dmol^−1^ to −55 deg cm^2^ dmol^−1^ is observed with increasing concentrations up to 300 g/L which demonstrates the disorderedness in the peptide. Also, a minimal shift in wavelength from 198 nm to 194 nm is detected. Overall, no noticeable changes are observed in spike cytoplasmic region in presence of crowders (**Figure 14**). These results are indicative of absence of weaker hydrophobic forces and electrostatic interactions among residues. These observations clearly explain that the spike endodomain or cytoplasmic domain is intrinsically disordered even under macromolecular crowding conditions.

**Figure 14:**
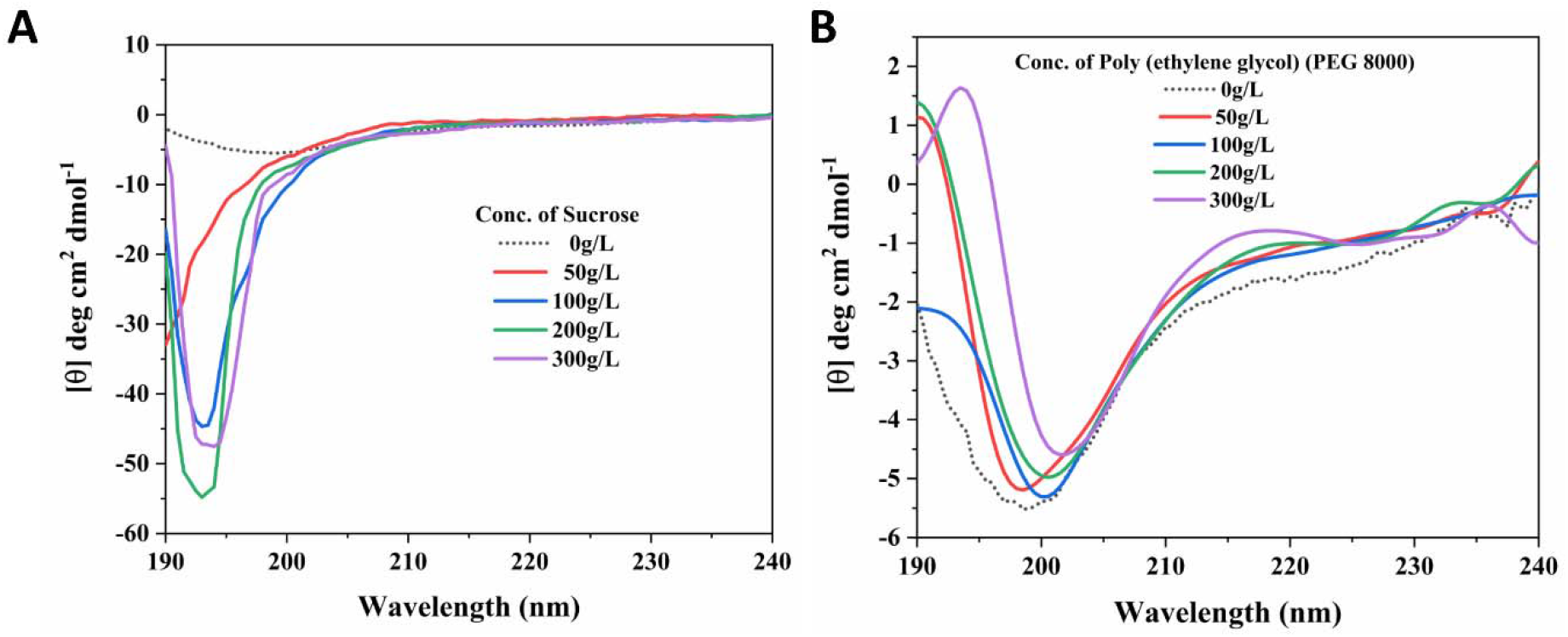
Effect of macromolecular crowding on spike cytoplasmic region (residues 1242-1273): **A.** and **B.** represent the far UV CD spectra spike cytoplasmic region in presence of sucrose and poly ethylene glycol (8000), respectively.

## Discussion

The existence of IDPs/IDPRs was established based on non-existing electron density of some regions in proteins during crystallography [12,46]. This unmapped electron density of characterized protein was later described as intrinsic disorderness of proteins. With increasingly published articles and revelation of presence of IDPs in frequently studied living systems, IDPs have been found prevalent in all three domains of life [46]. For a long period, the structure elucidation of transmembrane regions was not feasible due to their complex process of purification and crystallization [47]. It was also reported in a study on survey of disordered regions in eukaryotic proteins that the water-soluble cytoplasmic tails of membrane spanning proteins constitute intrinsic disorder [48]. These disordered regions are generally rich in certain residues such as proline, glycine, lysine and serine, and contains hydrophobic residues to form a core while folding [49]. The presence of disordered regions in proteins makes crystallization process an unsuccessful task and compel to produce a truncated structure.

In past decade, several reports of IDPs and IDRs in virus proteomes have exposed the “disordered” side of viral systems. It has answered some intriguing questions related to numerous protein-protein interactions led by viral proteins with host proteins required for hijacking the entire host cell [50]. Our detailed investigation on disordered regions in SARS-CoV and SARS-CoV-2 proteomes showed the presence of large flexible region in various proteins like nucleocapsid, ORF6, and Nsp8 proteins. The report also revealed the presence of intrinsic disorder in N- and C-terminal tails of most of the proteins [7]. It also exposed that the C-terminal region of spike protein of SARS-CoV-2 contain disordered regions. Recently, we have shown that the C-terminal regions of SARS-CoV-2 NSP1 and Envelope proteins are disordered in isolation [27,37].

Due to their dynamic nature, IDPs cannot be crystallized or frozen to be detected using such advanced structural biology techniques. X-ray crystallography and cryo-electron microscopy-based structures of SARS-CoV-2 spike proteins on PDB show missing electron density in transmembrane and cytosolic regions of C-terminal. Some of the structures of spike protein mentioned here with PDB IDs – 6VXX, 6ZGH, 6ZGG, 6XM3 and 6XM4, have reported the C-terminal tail as either unmodeled or as an artefact. Moreover, the consensus prediction of disorderness by MobiDB has also supported this observation by showing the missing residues consensus of region 1147-1273 which include transmembrane as well as cytosolic regions. The cytoplasmic domain is known to possess the localization signals for ER or endoplasmic reticulum-Golgi intermediate compartment in SARS-CoV as well as in other coronaviruses [4–6]. Notably, spike C-terminal tail contains multiple cysteine residues which may have implications in protein-protein interactions. In SARS-CoV, this cysteine-rich C-terminal domain of spike is responsible for interaction with M protein as a mutation in this domain obstructed their interaction [10,51].

Our study on the structural dynamics of SARS-CoV-2 spike cytoplasmic domain (residues 1242-1273) demonstrates it to be a disordered region. Based on the outcomes of two forcefields, OPLS 2005 and CHARMM36m, spike cytosolic region remains majorly unstructured. Additionally, in another simulation run of 1 μs, the cytoplasmic region with residues 1235-1273 have also shown a large part to be disordered and a small beta strand in few frames. As observed, the residues _**1257**_KFD_**1259**_ have shown propensity to form beta strands in simulations. Nevertheless, in REMD simulations, it adopted β-sheets at rising temperatures with time demonstrating its gain-of-structure property. However, we also tried to get the synthesized peptide of residues 1235-1273 of spike but due to multiple cysteine residues it was not feasible.

Further, it was of utmost importance to validate MD simulation outcomes using experimental techniques. The water-soluble peptide of spike residues 1242-1273 at 25 μM concentration exhibits a prominent negative peak at approximately 198 nm in far-UV CD spectra which defines the unstructured nature of a protein. Further, in presence of helix inducer solvent, TFE, the peptide adopts helical structure. However, SDS micelles in surroundings of peptide generates little changes in the peptide structure which may signify its inability to gain fold. In presence of crowding agents like sucrose and PEG (8000), conservation of disordered structure indicates that no -intra chain forces are acting in between the residues. Based on this combination of facts, we have interpreted that spike C-terminal cytosolic tail (residues 1242-1273) as an intrinsically disordered region.

## Conclusion

The cytoplasmic region of spike glycoprotein of SARS-CoV-2 has not been studied yet. Given its extreme importance in functioning of spike protein, the structure and its dynamics has been investigated here. The advancement in computational powers and excessive improvements in forcefields have empowered structural biology. Newly developed algorithms and their userfriendly approach allow correlating the outcomes with experimental observations. In this article, we have identified the transmembrane region in spike protein by employing distinguished web predictors. This cleared the composition of amino acids forming cytoplasmic domain. Further, the secondary structure and disorder predisposition analysis demonstrated it to be highly disordered. We have demonstrated the structural conformation of cytoplasmic domain (1242-1273 residues) of spike protein at a microsecond timescale using computational simulations. As revealed, this domain is purely unstructured or disordered after one microsecond and have not gained any structural conformation throughout the simulation period. Experimental outcomes also confirm the intrinsic disordered state of cytoplasmic domain of spike. The intrinsic disordered nature of peptide is shown in presence of macromolecular crowders. Based on our previous study [7], cytoplasmic tail of spike glycoprotein has molecular recognition features therein which needs to be explored further. In this study, the multiple conformations during the simulation process adds up to even more interesting speculations.

## Acknowledgements

All the authors would like to thank IIT Mandi for the infrastructure. RG is thankful to IYBA award from DBT, Government of India (BT/11/IYBA/2018/06), MHRD-SPARC (SPARC/2018-2019/P37/SL), and Science and Engineering Research Board (SERB), India (Grant Number: CRG/2019/005603). TB is thankful to DST for her INSPIRE fellowship. NG acknowledges Seed grant IOE, from Banaras Hindu University.

## Author contribution

RG, NG: Conception, design & study supervision. PK and TB: acquisition and interpretation of data. PK, TB, and RG: contributed to paper writing.

