## Supplementary File 1 for "Microsecond simulations and CD spectroscopy reveals the intrinsically disordered nature of SARS-CoV-2 Spike-C-terminal cytoplasmic tail (residues 1242-1273) in isolation"

**Supplementary Information**

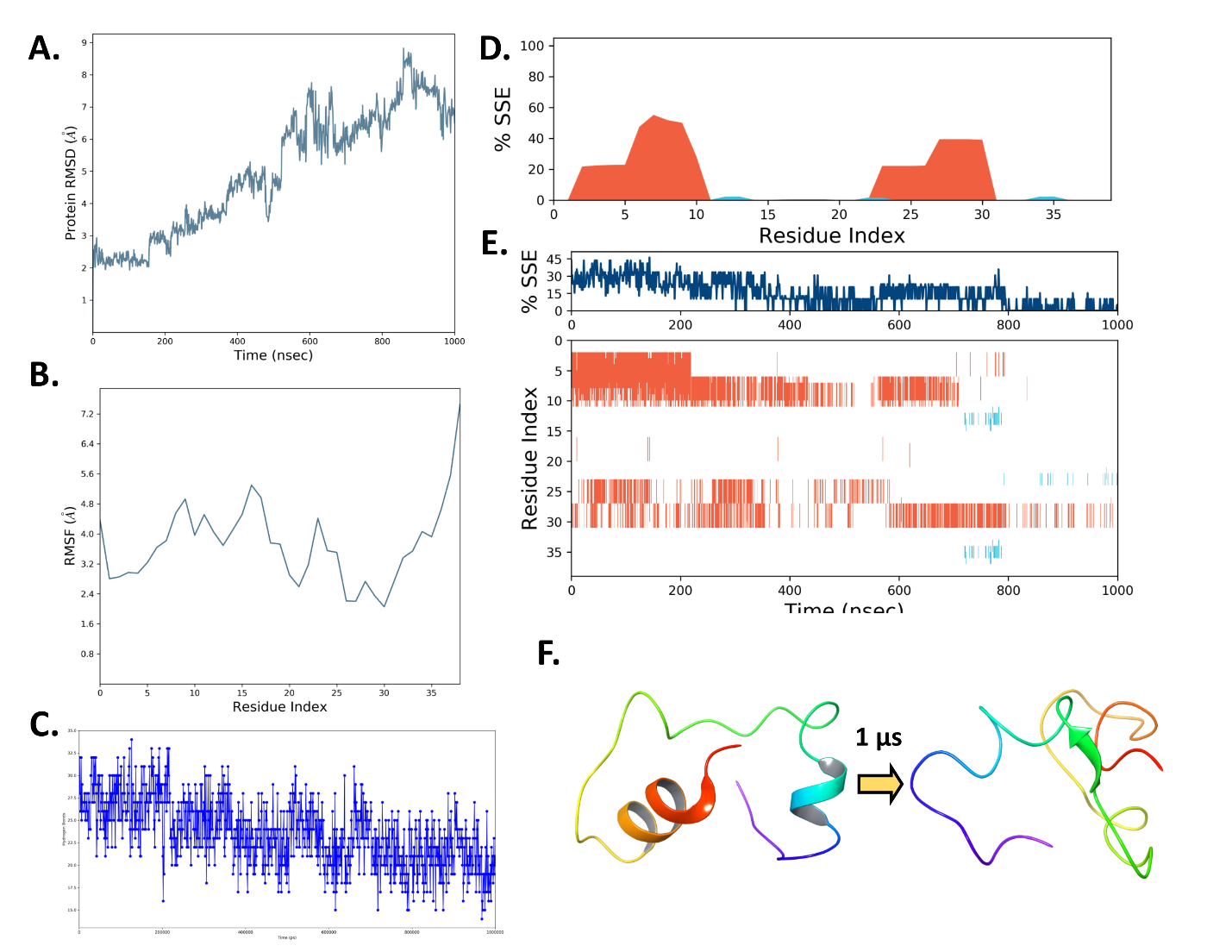

**Supplementary figure 1: One microsecond MD Simulation analysis of spike C-terminal cytoplasmic domain (residues 1235-1273): A.** Root mean square deviation (RMSD), **B.** Root mean square fluctuation (RMSF), **C.** Hydrogen bonds of protein, **D.** Secondary structure element (SSE) of residues, **E.** Timeline representation of secondary structure content during 1µs simulation time, and **F.** Representation of model to last frame after 1µs.

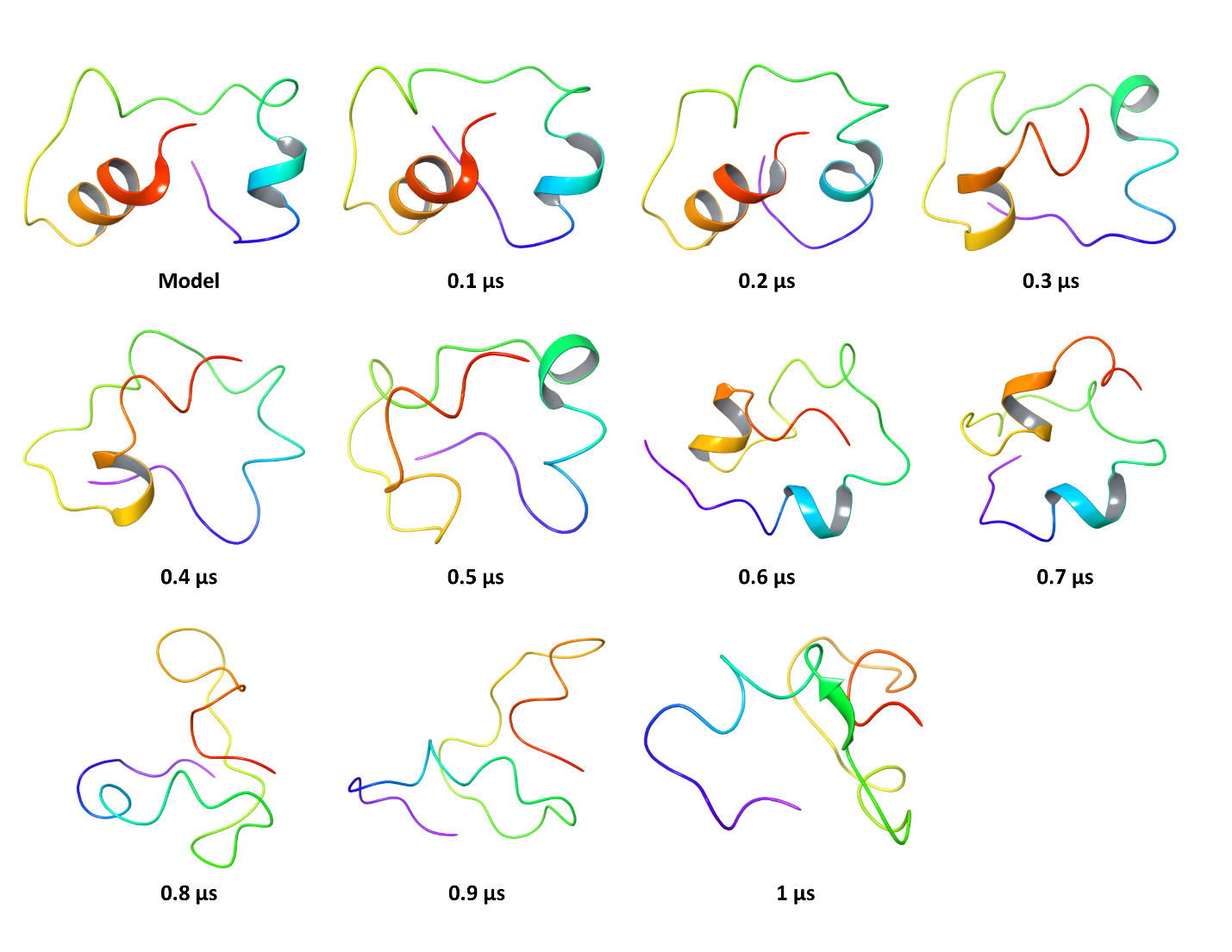

**Supplementary Figure 2: MD simulation of cytoplasmic region (1235-1273) with OPLS 2005 forcefield:** Snapshots at every 100 ns of 1 µs simulation trajectory.

**Supplementary Movie 1:** Desmond (OPLS 2005 forcefield) trajectory movie upto 1 µs simulation of Spike cytoplasmic region (residues 1242-1273).
